# Post-exercise cold-water immersion increases Na^+^,K^+^-ATPase α_2_-isoform mRNA content in parallel with elevated Sp1 expression in human skeletal muscle

**DOI:** 10.1101/151100

**Authors:** Danny Christiansen, Robyn M. Murphy, James R. Broatch, Jens Bangsbo, Michael J. McKenna, Jujiao Kuang, David J. Bishop

**Author notes:** Corresponding Author: David J. Bishop, Institute of Sport, Exercise and Active Living (ISEAL), College of Sport and Exercise Science, Victoria University, Footscray Park Campus, Building P, Cnr. Ballarat Rd/Geelong Rd, Footscray, Victoria 3011, Melbourne, Australia., Fax: n/a. **Abbreviations:** β2; β2M, β2-microglobulin; COLD, cold-water immersion treatment; CON, control treatment; Ct, cycle threshold; CV, coefficient of variation; CWI, cold-water immersion; EDL, extensor digitorum longus; FXYD1, phospholemman isoform 1; HIF-1α, hypoxia-inducible factor 1α; GAPDH, glyceraldehyde 3-phosphate dehydrogenase; GXT, graded exercise test; K^+^, potassium ion; mRNA, messenger RNA; Na^+^, sodium ion; NKA, Na^+^,K^+^-ATPase; PCR, polymerase chain reaction; RT, reverse transcription; ROS, reactive oxygen species; Sp1, specificity protein 1; Sp3, specificity protein 3; TBP, TATA-binding protein; VO2peak, maximum oxygen uptake.

## Abstract

We investigated the effect of a session of sprint-interval exercise on the mRNA content of NKA isoforms (α_1-3_, β_1-3_) and FXYD1 in human skeletal muscle. To explore some of the cellular stressors involved in this regulation, we evaluated the association between these mRNA responses and those of the transcription factors Sp1, Sp3 and HIF-1α. Given cold exposure perturbs muscle redox homeostasis, which may be one mechanism important for increases in NKA-isoform mRNA, we also explored the effect of post-exercise cold-water immersion (CWI) on the mRNA responses. Muscle was sampled from nineteen men before (Pre) and after (+0h, +3h) exercise plus passive rest (CON, n=10) or CWI (10°C; COLD, n=9). In COLD, exercise increased NKAα_2_ and Sp1 mRNA (+0h, p<0.05). These genes remained unchanged in CON (p>0.05). In both conditions, exercise increased NKAα_1_, NKAβ_3_ and HIF-1α mRNA (+3h; p <0.05), decreased NKAβ_2_ mRNA (+3h; p<0.05), whereas NKAα_3_, NKAβ_1_, FXYD1 and Sp3 mRNA remained unchanged (p>0.05). These human findings highlight 1) sprint-interval exercise increases the mRNA content of NKA α_1_ and β_3_, and decreases that of NKA β_2_, which may relate, in part, to exercise-induced muscle hypoxia, and 2) post-exercise CWI augments NKAα_2_ mRNA, which may be associated with promoted Sp1 activation.

## Introduction

It is well-established that the contractile function of skeletal muscle is limited by the capacity of the active transport system, the Na^+^,K^+^-ATPase (NKA), to counterbalance the loss of potassium (K^+^) ions and the gain in sodium (Na^+^) ions within the muscle cells. The NKA is a heterotrimeric complex composed of a catalytic α, a regulatory β, and an accessory (phospholemman; FXYD) subunit (1). Different isoforms of each of these subunits have been identified in human skeletal muscle (α_1-3_, β_1-3_ and FXYD1) (2, 3). As the expression of NKA isoforms determines, in part, the difference in the capacity for Na^+^/K^+^ transport between muscles of different fibre type (4), identifying novel strategies to enhance their expression in human muscle is of fundamental importance in the context of both disease prevention (5) and sport performance (6).

One proposed mechanism for the upregulation of muscle NKA-isoform protein abundance with exercise training is the transient bursts in isoform mRNA content that accompany repeated exercise sessions, as demonstrated for mitochondrial proteins (7). This is supported by evidence from a study of rats (8), and studies of mRNA changes to a one-off exercise session in humans (2, 9-12). However, it remains to be studied how the mRNA content of NKA isoforms is affected by repeated-sprint exercise (i.e. maximal-intensity bouts < 1 min in duration). Furthermore, it is unknown whether FXYD1 is regulated at the mRNA level in human muscle by contractile activity.

In *in vitro* cell culture models, the GC box-binding transcription factors, specificity protein 1 and 3 (Sp1 and Sp3), have been shown to mediate NKA (α_1_ and β_1_) transcription (13, 14). Activation of these factors by transcriptional or post-transcriptional modification occurs in response to ROS-mediated oxidative stress (15) or perturbations in cellular redox state (13, 14). In contrast, the transcription factor, hypoxia-inducible factor 1α (HIF-1α), is upregulated transcriptionally under chronic exposure to cellular hypoxia (16). Thus, insights into which cellular signals are important for contraction-stimulated NKA adaptation in humans *in vivo* might be gained from measuring changes in Sp1, Sp3 and HIF-1α mRNA concomitantly with NKA mRNA content following the same exercise session.

There is great interest in the use of cold-water immersion (CWI) to optimize muscle recovery, and how it may affect adaptations to exercise training (17-19). In humans, cold exposure has been shown to increase the systemic level of norepinephrine (20), which has been linked to greater oxidative stress (21). In many cell types, cold exposure has also been reported to perturb redox homeostasis and increase the production of reactive oxygen species (ROS) (22). Given these cellular stressors could be important for the isoform-dependent modulation of NKA mRNA content in human muscle (23), post-exercise CWI could serve as a potent stimulus to modify NKA-isoform expression in this tissue. However, no study has investigated the effect of post-exercise CWI on muscle NKA mRNA content in humans.

The aims of the current study was to examine in human skeletal muscle 1) the effect of a session of repeated-sprint exercise on the mRNA content of NKA isoforms (α_1-3_, β_1-3_) and FXYD1, 2) if changes in the content of these isoforms are associated with modulation of the expression of genes encoding the transcription factors Sp1, Sp3 and HIF-1α, and 3) the effect of post-exercise CWI on these mRNA responses. In this acute setting, these examinations will inform about the potential of whether transcriptional regulation of these proteins is evident.

## Materials and Methods

### Ethical Approval

This study received ethics approval by the Human Research Ethics Committee of Victoria University, Australia (HRE12-335) and was performed in accordance with the latest guidelines in the Declaration of Helsinki. The participants were informed in writing and orally about the procedures, potential risks and benefits of the study prior to providing oral and written consent to participate.

### Participants

Nineteen healthy males with an age, body mass, height and peak oxygen uptake (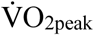) of (mean ± SD) 24 ± 6 y, 79.5 ± 10.8 kg, 180.5 ± 10.0 cm and 44.6 ± 5.8 mL·kg^-1^·min^-1^, respectively, were recruited for this study. They were healthy, physically active several days per week, and non-smokers. In addition, they were free of medicine and anti-inflammatory drugs during the study. This study was part of a larger research project investigating the effect of post-exercise cold-water immersion. Protein data for this project is published elsewhere (Christiansen et al. 2017, in review, The Journal of Physiology).

### Experimental design

Before the first biopsy session, participants were familiarized with the exercise protocol and recovery treatments on one occasion. In this period, they also performed a graded exercise test (GXT) to volitional fatigue on an electromagnetically-braked cycle ergometer (Lode, Groningen, The Netherlands) to determine their 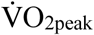. These visits were separated by at least 24 h and were completed at least 3 days prior to the biopsy session. This study utilized a parallel, two-group design, which is illustrated in Fig. 1. After being matched on their 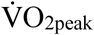, participants were randomly assigned by a random number generator (Microsoft Excel, MS Office 2013), in a counter-balanced fashion, to one of two recovery treatments that concluded a repeated-sprint exercise session: Cold-water immersion (COLD, *n* = 9) or non-immersion rest at room temperature (CON, *n* = 10). A muscle biopsy was obtained at rest before exercise (Pre), 2 min post (+0h), and 3 h after (+3h), the allocated treatment to quantify mRNA expression.

**Figure 1.**
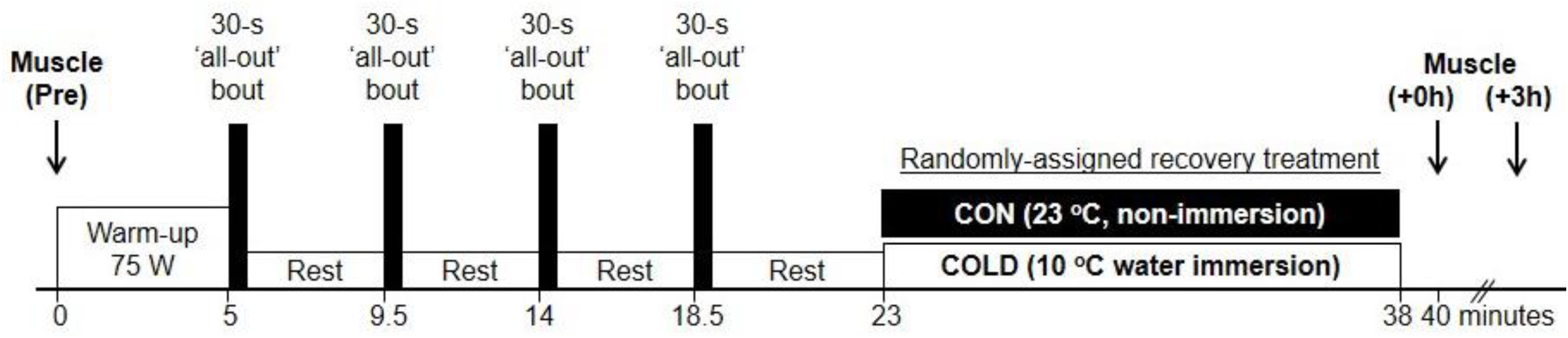
A time-aligned representation of the experimental setup. Muscle was sampled at rest before exercise (Pre), 2 min post (+0h), and 3 h after (+3h), 15-min of passive rest at room temperature (CON group) or cold-water immersion up to the umbilicus (~10°C; COLD group) that followed an intermittent sprint exercise session.

### Repeated-sprint protocol

All experimental sessions took place in the Exercise Physiology Laboratory at the Institute of Sport, Exercise and Active Living (ISEAL), Victoria University (Melbourne, Australia) under standard laboratory conditions (~23°C, ~35% relative humidity). The participants performed the repeated-sprint session on the same electrically-braked cycle ergometer as used during the prior visits. The exercise protocol consisted of a 5-min warm-up at a constant absolute intensity (75 W), followed by four 30-s maximal-intensity (‘all-out’) sprints at a constant relative flywheel resistance of 7.5 % of body mass, interspersed by 4 min of passive recovery, in which the participants remained seated with their legs resting in the pedals. Each sprint was executed from a flying start at ~120 rpm prepared by one of the investigators by manual rotation of the flywheel. The participants remained seated in the saddle during the entire session. One investigator provided verbal encouragement to the participants throughout each sprint.

### Recovery treatments

The participants commenced their assigned 15-min recovery treatment five minutes after termination of exercise. The recovery treatments were either rest in a seated posture with the legs fully extended on a laboratory bed at room temperature (~23°C, CON), or 10°C water immersion up to the umbilicus in the same position in an inflatable bath (COLD; iBody, iCool Sport, Miami QLD, Australia). A cooling unit (Dual Temp Unit, iCool Sport, Miami QLD, Australia) ensured a constant water temperature and water agitation.

### Muscle sampling

Each muscle biopsy was obtained from the *vastus lateralis* muscle of the participants’ right leg using a 5-mm Bergström needle with suction. Prior to sampling, a small incision was made through the skin and fascia under local anesthesia (5 mL, 1% Xylocaine). Separate incisions were made for the biopsies, which were separated by ~1-2 cm to minimize interference of prior muscle sampling on the mRNA response (24). Participants rested on a laboratory bed during each sampling procedure and between the second and third sampling time point. Immediately post sampling, the biopsies were rapidly blotted on filter paper to remove excessive blood, and instantly frozen in liquid nitrogen. The samples were stored at −80°C until subsequent analysis. The incisions were covered with sterile Band-Aid strips and a waterproof Tegaderm film dressing (3M, North Ryde, Australia).

### RNA isolation

From each biopsy, ~25 mg w.w. muscle was added to 1 g zirconia/silica beds (1.0 mm, Daintree Scientific, Tasmania, Australia), and homogenized in 800 μL TRIzol reagent (Invitrogen, Carlsbad, CA) using an electronic homogenizer (FastPrep FP120 Homogenizer, Thermo Savant). After centrifugation (15 min at ~12.280 *g*), cell debris was removed, and the supernatant added to 250 μL chloroform (Sigma Aldrich, St. Louis, MO) and centrifuged (15 min at ~12.280 *g*) at 4°C. RNA was precipitated by aspirating the superior phase into a new microfuge tube containing 400 μL 2-isopropanol alcohol (Sigma-Aldrich, St Louis, MO) and 10 μL of 5 M NaCl. Following storage at −20°C overnight, samples were centrifuged (20 min at ~12.280 *g*) at 4°C, after which the isopropanol was aspirated. The RNA pellet was rinsed with 75% ethanol made from DEPC-treated H2O (Invitrogen Life Sciences), and centrifuged (~5890 *g* for 8 min) at 4°C. After aspirating the ethanol, the pellet was suspended in 5 μL of heated DEPC-treated H2O. RNA concentration and purity was determined spectrophotometrically (NanoDrop 2000, Thermo Fisher Scientific, Wilmington, DE) at 260 and 280 nm. The RNA yield was 1252 ± 467 ng·μL^-1^ and the ratio of absorbance (260nm/280nm) was 1.78 ± 0.11. RNA was stored at -80°C until reverse transcription.

### Reverse transcription

For each sample, 1 μg of RNA was transcribed into cDNA on a thermal cycler (S1000™ Thermal Cycler, Bio-Rad, Hercules, CA) using the iScript™ cDNA Synthesis Kit (Bio-Rad, Hercules, CA) and the following incubation profile: 5 min at 25 °C, 30 min at 42 °C and 5 min at 85 °C. The transcription was performed with random hexamers and oligo dTs in accordance with the manufacturer’s instructions. cDNA was stored at - 20°C until subsequent analysis.

### Primers

Primers were designed using Primer BLAST (National Centre for Biotechnology Information) and are shown in Table 1. To improve the construct validity of product amplification by real-time PCR, primers were designed to target a region on the genes encoding for most splice variants and to ensure sequence homology for the target gene only. Furthermore, primers fulfilled 3’end self-complementarity < 3.00, span of an exon-exon junction, and PCR product size < 150 bp. The difference in maximum melting temperature between forward and reverse primers was < 2 °C. The β2M gene primer set was adopted from Vandesompele, et al. (25). Primer validation was performed in two steps. First, its optimal annealing temperature (i.e. resulting in the highest yield with no non-specific amplification) was determined using gradient PCR, and the result verified by agarose gel (2 %) electrophoresis. If primers were successful, their efficiency was evaluated by real-time PCR of a 10 × cDNA dilution series. Primer efficiency was calculated from the slope of the standard curve generated from the log-transformed cDNA dilutions and corresponding C_t_. From these results, each reaction was designed to yield a C_t_ within the linear range of detection by adjusting input cDNA concentration for every gene quantified.

**Table 1.**
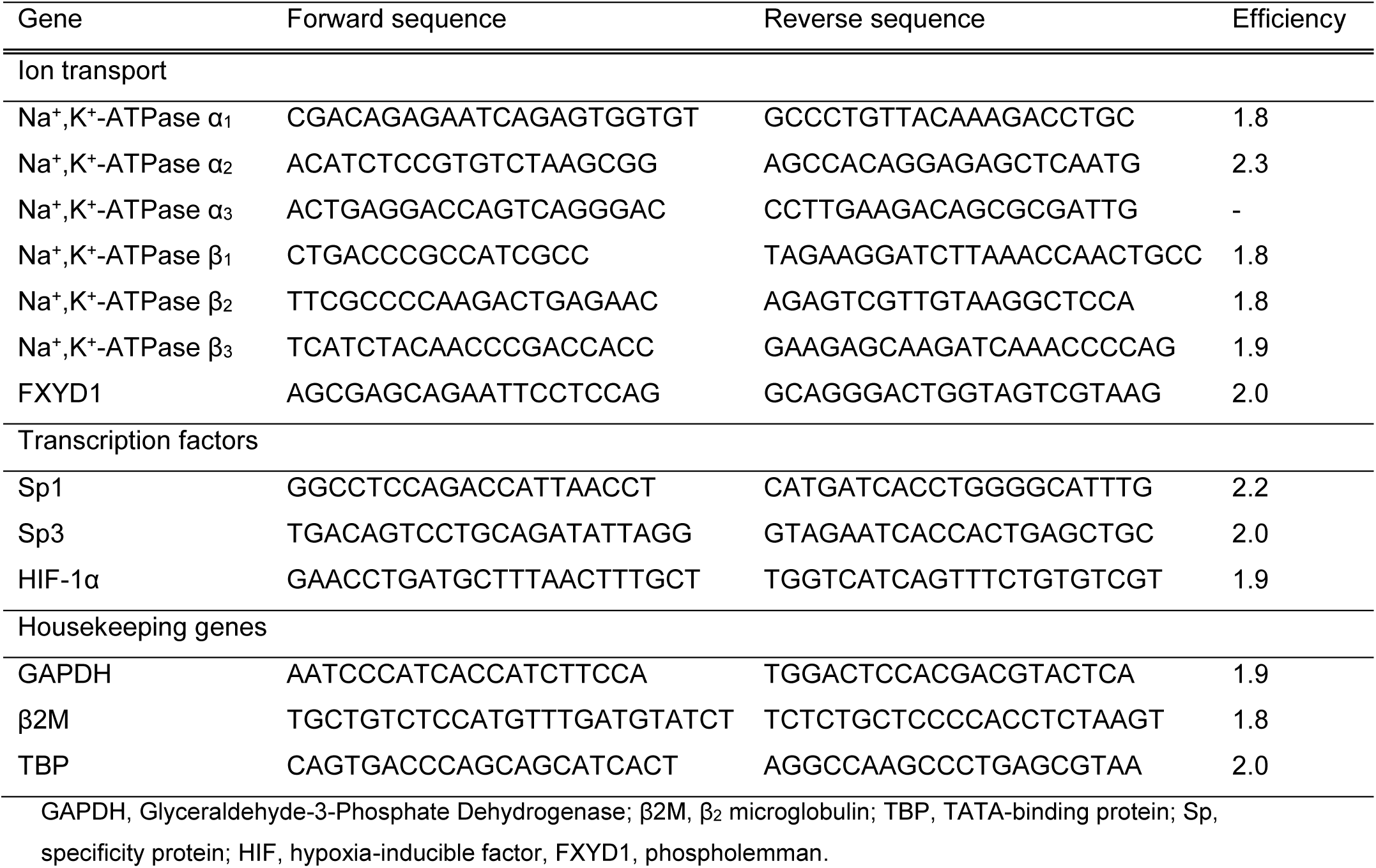
Forward and reverse primer sequences used to determine gene expression of Na+,K+-ATPase isoforms, transcription factors and reference genes, and their amplification efficiency during real-time PCR

### Real-time PCR and mRNA data treatment

mRNA expression was quantified by real-time PCR on a Mastercycler RealPlex 2 (Eppendorf, Hamburg, Germany). Each reaction (10 μL) was composed of 4 μL of diluted cDNA, 5 μL of 2 × concentrated iTaq universal SYBR Green supermix (Bio-Rad, Hercules, CA), with SYBR Green I as the fluorescent agent, 0.6 μL of primer diluted in DEPC-treated H_2_O and 0.4 μL DEPC-treated H2O. Reactions were denatured at 95 °C for 3 min to activate the enzyme before PCR cycling. 40 PCR cycles were executed by heating to 95 °C for 15 s, followed by 60 °C for 60 s. Samples and template-free controls were loaded in duplicate on the same plate and samples from the same subject from all time points were on the same plate. Reactions were prepared using an automated pipetting system (epMotioon 5070, Eppendorf, Hamburg, Germany). The expression of target genes was normalized to that of three housekeeping genes using the 2^-ΔΔCt^ method (26). This correction has been shown to yield reliable and valid mRNA data (25). Housekeeping genes used were glyceraldehyde 3-phosphate dehydrogenase (GAPDH), TATA-binding protein (TBP) and β2 microglobulin (β2M) as their mRNA expression remained unchanged with our exercise protocol (p > 0.05, using 2^-Ct^; data not shown). The mean (± SD) intra-assay and inter-assay coefficient of variation for the investigated genes was 5.8 % ± 2.6 and 3.0 % ± 1.1, respectively, and are shown for each gene in Table 2. Prior to statistical analysis, an iterative elimination of outliers was performed using the following criteria: An arbitrary expression at pre deviating >3 fold from the group mean. Four outliers were identified: one in CON for α_1_ (5.3 fold) and one in COLD for α_1_, α_2_ and β_1_ (4.7, 3.1 and 3.5 fold, respectively). Thus, the sample size in CON and COLD, respectively, was: *n* = 9 and 8 for α_1_, *n* = 10 and 8 for α_2_ and β_1_, and *n* = 10 and 9 for the other genes.

**Table 2:**
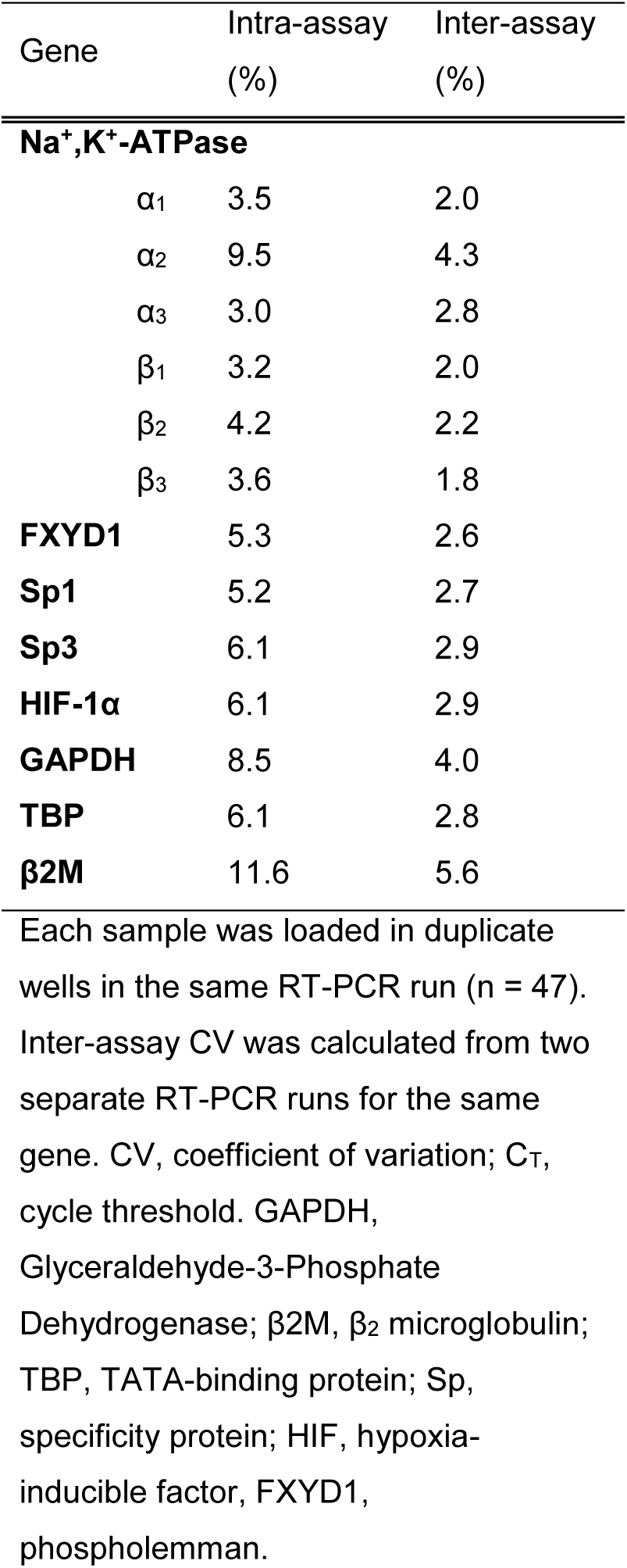
Intra- and inter-assay variability of raw C_T_ values for mRNA expression determined with RT-PCR.

### Statistical analysis

Prior to parametric tests, data was assessed for normality using the Shapiro-Wilk test. An appropriate transformation was used, if required, to ensure a normal distribution. A two-way repeated-measures (RM) ANOVA was used to test the effect of group (COLD, CON) and time (Pre, +0h, +3h) using the 2^-ΔΔCt^ expression data. Where applicable, multiple pairwise, *post hoc*, comparisons used the Tukey test. Cohen’s conventions were adopted for interpretation of effect size (*d*), where <0.2, 0.2-0.5, >0.5-0.8 and >0.8 were considered as trivial, small, moderate and large, respectively (27). Data are reported as geometric mean ± 95% confidence intervals (CI95) in figures. Note that the mRNA expression at Pre is not equal to 1.0 due to the nature of using geometric means. F statistic (F), and *d* +/- CI95 are shown for time effects and group interactions. The α-level was set at p ≤ 0.05. Statistical analyses were performed in Sigma Plot (Version 11.0, Systat Software, CA).

## Results

In figures, individual data are displayed on the left with each symbol representing the same participant for all genes, and geometric means ± CI95 are shown on the right. Fold-changes are reported relative to the participants’ geometric mean at Pre in CON.

### Na^+^,K^+^-ATPase α isoforms

In both CON and COLD, α_1_ mRNA increased from Pre to +3h (p < 0.001; *d* = 1.28±0.50 and 1.29±0.63, respectively), with no difference between groups (group × time interaction: F = 0.07; p = 0.802; Fig. 2A). In COLD, α_2_ mRNA increased from Pre to +0h (p = 0.013; *d* = 1.2±0.9). In contrast, it remained unchanged in CON (p = 0.862; *d* = 0.20±0.31; Fig. 2B). This change in COLD was higher than in CON (group × time interaction: 1.42 ± 1.32 fold; p < 0.001, *d* = 1.23±0.63; Fig. 2B). In both groups, α_3_ mRNA content remained unchanged following exercise (F = 0.68; p = 0.513; pooled *d* = 0.23±0.25), with no difference between groups (group × time interaction: F = 0.56; p = 0.578; Fig. 2C).

**Figure 2.**
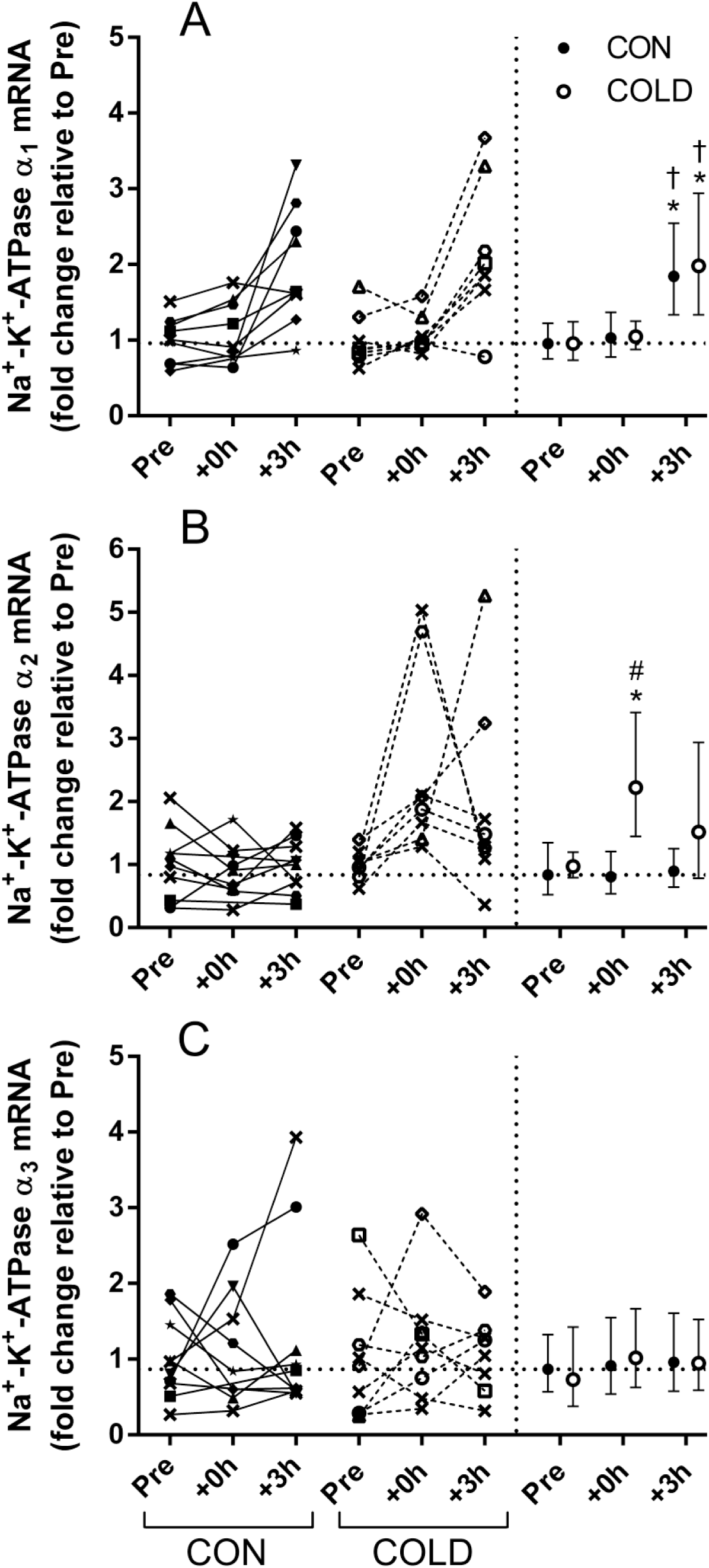
Effect of a single session of sprint-interval exercise with (COLD) or without (CON) post-exercise cold-water immersion on Na^+^,K^+^-ATPase α-isoform gene expression. A) α1, B) α2 and C) α3 mRNA expression. Muscle was sampled at rest before exercise (Pre), 2 min post (+0h) and 3 h after (+3h) the assigned 15-min post-exercise recovery treatment. The recovery treatment commenced 4 min after termination of exercise. Individual values (left) and geometric mean ± 95% confidence intervals (right) are displayed on each graph for CON (● closed symbols) and COLD (○ open symbols). Each symbol represents one participant (left) and is the same for gene and protein data (Fig. 8). Symbol **x** represents participants, from which no muscle sample was left to determine protein abundance. The horizontal, dotted line represents the geometric mean expression at Pre in CON. *p < 0.05, different from Pre within group; †p < 0.05, different from +0h within group; #p < 0.05, Pre to +0h change and time point different from CON.

### Na^+^,K^+^-ATPase isoforms

In both groups, β_1_ mRNA remained unchanged following exercise (F = 0.36; p = 0.700; pooled *d* =0.29±0.31), with no difference between groups (group × time interaction: F = 0.05; p = 0.949; Fig. 3A). In both CON and COLD, β_2_ mRNA content decreased from Pre to +3h (p = 0.002, *d* = 0.86±0.21; and p =0.008, *d* = 0.82±0.27, respectively), with no difference between groups (group × time interaction: F = 0.02; p = 0.980; Fig. 3B). In both CON and COLD, β_3_ mRNA increased from Pre to +3h (p < 0.001, *d* =1.23±0.57; and p < 0.001, *d* = 1.39±0.86, respectively), with no difference between groups (group × time interaction: F = 0.52; p = 0.598, Fig. 3C).

**Figure 3.**
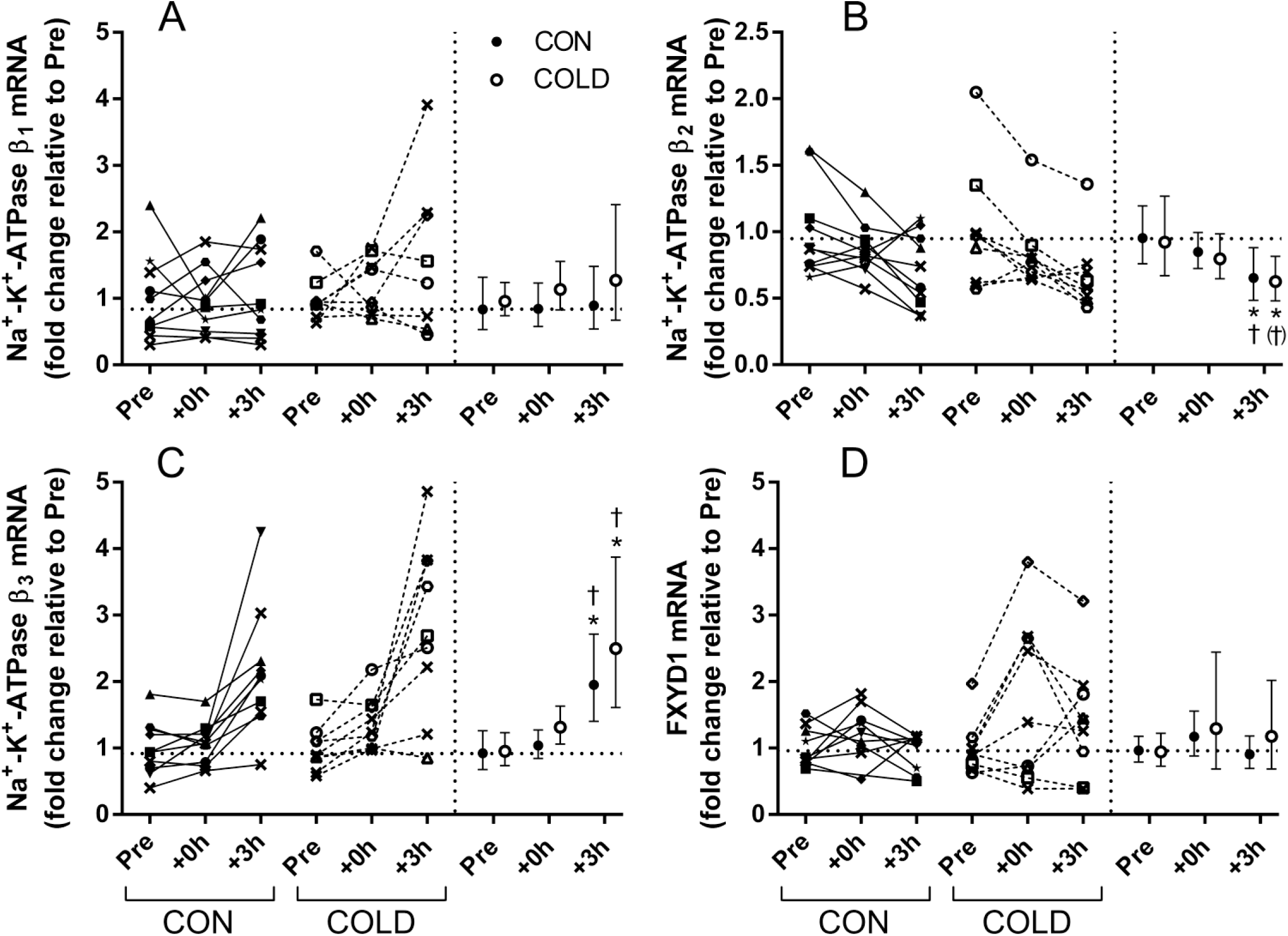
Effect of a single session of sprint-interval exercise with (COLD) or without (CON) post-exercise cold-water immersion on Na+,K+-ATPase β-isoform and FXYD1 gene expression. A) β_1_, B) β_2_, C) β_3_, and D) FXYD1 mRNA expression. Muscle was sampled at rest before exercise (Pre), 2 min post (+0h) and 3 h after (+3h) the assigned post-exercise recovery treatment. The recovery treatment was employed from 4 to 19 min after termination of exercise. Individual values (left) and geometric mean ± 95% confidence intervals (right) are displayed on each graph for CON (● closed symbols) and COLD (○ open symbols). Each symbol represents one participant (left) and is the same for gene and protein data (Fig. 9). Symbol **x** represents participants, from which no muscle sample was left to determne protein abundance. The horizontal, dotted line represents the geometric mean expression at Pre in CON. *p < 0.05, different from Pre within group; †p < 0.05, different from +0h within group; (†)p = 0.064, different from +0h within group.

### Phospholemman (FXYD1)

In both groups, FXYD1 mRNA remained unchanged following exercise (F = 0.68; p = 0.512; pooled *d* =0.45±0.51), with no difference between groups (group × time interaction: F = 0.05; p = 0.952; Fig. 3D).

### Transcription factors

The change in Sp1 mRNA from Pre to +0h in COLD was 0.67 ± 1.07 fold greater than in CON (p = 0.044; *d* = 0.63±0.51; Fig. 4A). Despite a moderate effect in COLD (*d* = 0.6), Sp1 mRNA was not significantly increased from Pre to +0h in both groups (CON: p = 0.154; *d* = 0.34±0.40; COLD: p = 0.094; *d* = 0.57±0.77). Sp3 mRNA remained unchanged following exercise in both groups (main effect for time: F =1.00; p = 0.378; pooled *d* = 0.09±0.21; Fig. 4B), with no difference between groups (group × time interaction: F = 0.33; p = 0.723). HIF-1α mRNA tended to increase from Pre to +3h in both groups (CON: p = 0.059; *d* = 0.93±0.58; COLD: p = 0.071; *d* = 0.62±0.67), with no difference between them (group × time interaction: F = 0.47; p = 0.631). Using the pooled data (both groups), HIF-1α mRNA increased from Pre to +3h (main effect for time: F = 5.99; p = 0.006; pooled *d* = 0.51±0.42; Fig. 4C).

**Figure 4.**
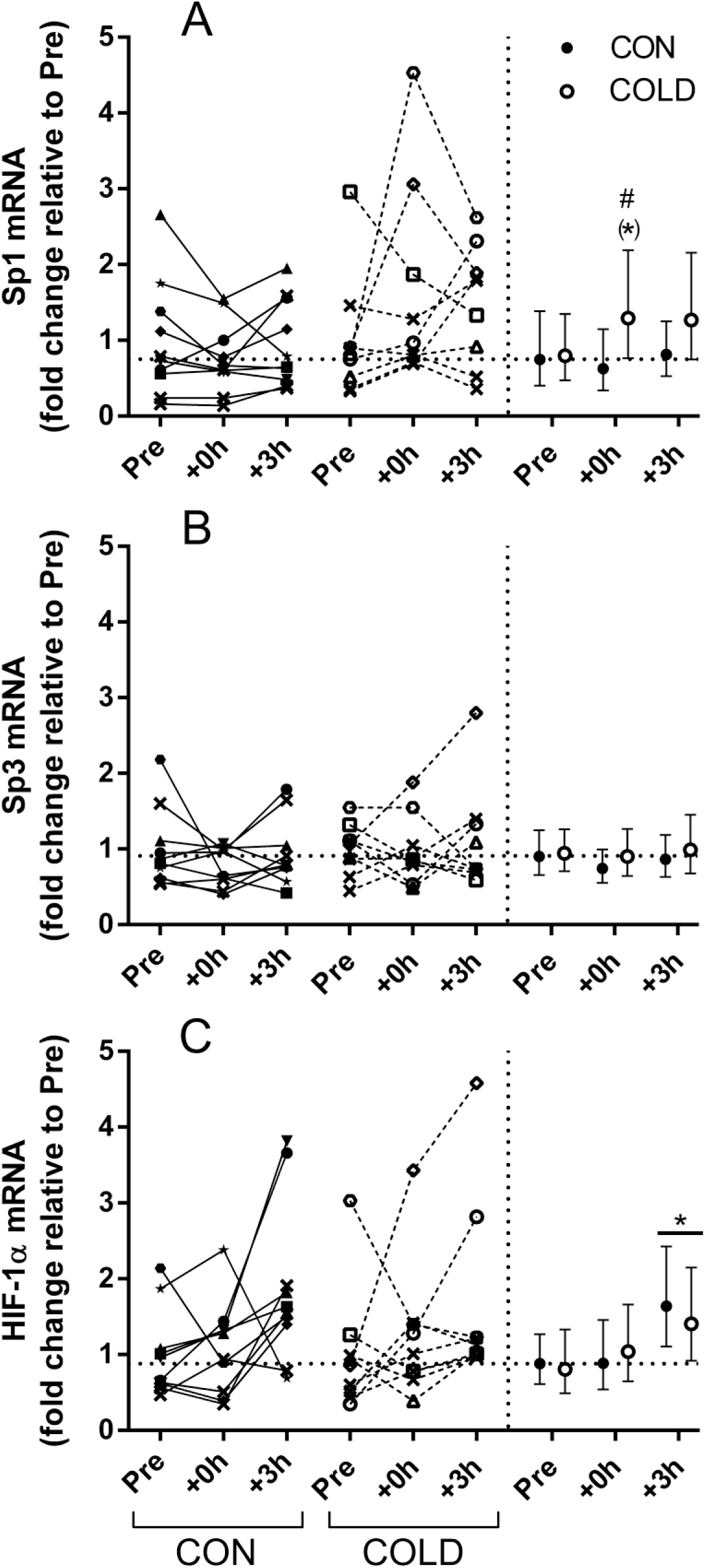
Effect of a single session of sprint-interval exercise with (COLD) or without (CON) post-exercise cold-water immersion on Sp1, Sp3 and HIF-1α gene expression. Individual values (left) and geometric mean ± 95% confidence intervals (right) are displayed on each graph. Representation of symbols as per Fig. 2 and 3. The horizontal, dotted line represents the geometric mean expression at Pre in CON. Muscle was sampled at rest before exercise (Pre), 2 min post (+0h) and 3 h after (+3h) the assigned post-exercise recovery treatment. *p < 0.05, different from Pre; #p < 0.05, Pre to +0h change different from CON; (*)p = 0.094, different from Pre within group.

## Discussion

This study yields the following main novel findings, which are summarized in figure 5: 1) A single session of repeated-sprint exercise increased the mRNA content of NKA α_1_ and β_3_, and decreased that of NKA β_2_, in temporal association with elevated HIF-1α mRNA content, whereas the repeated-sprint exercise session was without effect on NKA α_2_, α_3_, β_1_, FXYD1, and the transcription factors Sp1 and Sp3, 2) Post-exercise cold-water immersion (CWI) transiently increased the mRNA content of the NKA α_2_ isoform in parallel with elevated Sp1 expression. Given the positive association between Sp1 mRNA content and its DNA binding activity (28), our results support that the transcription and/or stabilization of the α_2_ mRNA transcript in human muscle may be promoted by Sp1 activation (13, 29) following a bout of post-exercise cold-water immersion.

**Figure 5.**
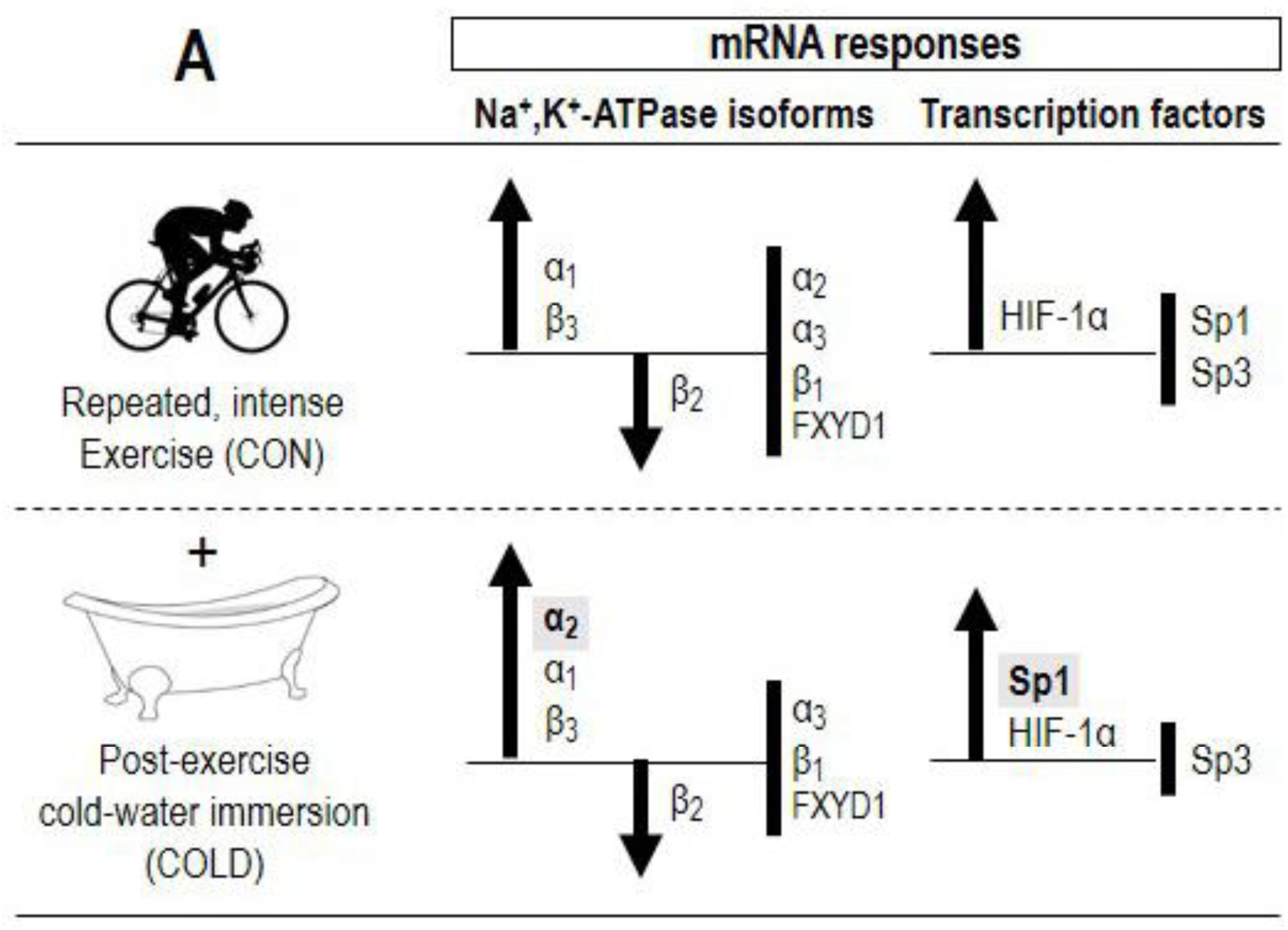
Summary of the key findings. Effect of a single session of sprint-interval exercise without (CON) or with post-exercise cold-water immersion (COLD) on the mRNA expression of Na^+^,K^+^-ATPase isoforms (α_1-3_ and β_1-3_), phospholemman (FXYD1) and redox-sensitive transcription factors (Sp1, Sp3 and HIF-1α). Note the selective increases in NKA α_2_ and Sp1 mRNA content with COLD. Bold vertical lines without arrow indicate NKA isoforms or transcription factors that remained unchanged with the given intervention.

### Regulation of the NKA α_1_ and β_3_ mRNA transcripts in human muscle by repeated-sprint exercise

A novel finding was that the NKA α_1_ mRNA content increased by 2 fold in human skeletal muscle after one session of repeated-sprint exercise. This is in agreement with the increase (2.5-3.8 fold) in muscle α_1_ mRNA in both untrained and trained men after a session of intense intermittent cycling (30, 31) or continuous, fatiguing knee-extensor exercise (2). Similar to the mRNA response for α_1_, β_3_ mRNA content increased by 2 fold, which is within the range of increases (1.9-3.1 fold) previously reported in recreationally active men after knee-extensor exercise (5 × 2-5 min at 56 W, 3 min rest; 10) or continuous cycling (~75 min at 60-85 % VO2peak; 32). The mRNA responses for α_1_ and β_3_ were unrelated to the induction of Sp1 after CWI (Fig. 4). Given the positive association between the mRNA content of Sp1 and its DNA binding activity (33), these results support that Sp1 is unlikely to activate, but may still take part in the regulation of, these NKA transcripts in human muscle. Accordingly, a dissociation of NKA α_1_ mRNA content with Sp1 DNA binding activity was shown in rat gastrocnemius muscle (28). In contrast, the induction of the α_1_ and β_3_ transcripts was temporally associated with an increase in HIF-1α mRNA, indicating cellular hypoxia may be one stressor underlying upregulation of these mRNA transcripts with repeated-sprint exercise in human muscle (16). Contrary to the present observation, β_3_ mRNA remained unaltered in other human exercise studies (2, 23, 30, 34). This controversy could be attributable, at least in part, to the highly-trained (VO_2peak_ ~62-66 mL· kg^-1^· min^-1^) and sex-heterogeneous cohorts utilized in some of the latter studies (2, 34), as these factors are likely to modulate β_3_ mRNA content in human muscle (35), whereas the early (+0 h) muscle sampling time point post exercise might have been influential in others (23, 30).

### Regulation of the NKA α_2_ mRNA transcript in human muscle by post-exercise cold-water immersion

Another novel finding was that post-exercise CWI augmented (2.2 fold) the α_2_ mRNA expression, whereas repeated-sprint exercise alone was without any effect. In humans, CWI (10 min at 8°C) has been reported to increase the systemic concentration of norepinephrine (20), and exacerbate perturbations in muscle convective oxygen delivery during exercise (36). In combination, these alterations could create a potent cellular environment for ROS formation (37). By intravenous infusion with the antioxidant N-acetylcysteine during contractions *in vivo*, Murphy, et al. (23) demonstrated that ROS may be involved in the regulation of α_2_ mRNA content in human skeletal muscle (23). Thus, one mechanism underlying the effect of CWI in the current study might be its ability to modulate myocytic redox state and, consequently, muscle ROS generation. To assess this possibility indirectly in humans *in vivo*, we compared the temporal patterns of changes in the mRNA content of NKA isoforms with those of the transcription factors Sp1, Sp3 and HIF-1α, as these are known to be modulated differently according to the fluctuations in cellular redox state (15, 16). Interestingly, the rise in α_2_ mRNA content promptly after termination of CWI coincided with increased Sp1 mRNA content (67 % vs. CON, moderate effect; Fig. 4). In cell culture models, Sp1 has been shown to regulate NKA mRNA transcription in a ROS-dependent manner (38), and the 5’-flanking region of the α_2_ gene contains multiple binding sites for Sp1 (29) (13). This, along with the present results, indicates that Sp1 could be important for the transcriptional activation or stabilization (or both) of the α_2_ gene in human skeletal muscle. This is partly supported by the observation that α_2_ expression was reduced concomitant with a decrease in Sp1 DNA binding activity in sedentary rats exposed to chronic intake of fat-rich meals (28). However, further experimental verification of our initial human findings is required (e.g. by use of a chromatin immunoprecipitation ChiP assay).

### Regulation of the NKA β_2_ mRNA transcript in human muscle by repeated-sprint exercise

We also report for the first time in humans that muscle β_2_ mRNA content decreased (to ~0.65 fold) after a single session of repeated-sprint exercise. This result is consistent with a downregulation (to ~0.4-fold of pre level) in the number of β_2_ mRNA transcripts in rat EDL muscle after electrical stimulations *in vivo* (20 Hz for 400 ms every 2 s for 40 min; 11). The decrease in β_2_ mRNA was paralleled by an increase in HIF-1α mRNA. As HIF-1α mRNA stability and transcription increases during cellular hypoxia (39), our results could indicate that β_2_ mRNA stability is adversely influenced by a hypoxic muscular environment. In contrast to our result, β_2_ mRNA increased (10, 11, 23), or remained unaltered (2, 9, 30, 34), in other human studies. This discrepancy may in part be explained by the different exercise modes used, since sprint-interval and endurance training induce opposite changes in β_2_ mRNA content in rat muscles (8). In that same study, mRNA responses to the first training session was dissociated from protein changes after both 3 days and 3 weeks of training. It is therefore possible that our decrease in β_2_ mRNA content represented a transient state of mRNA decay with negligible impact on mRNA translation and/or protein synthesis. The fact that the first step in the deadenylation-dependent mRNA decay pathway is reversible (40) supports this proposition. Future work would need to examine the association between mRNA responses to exercise sessions with those of protein to weeks of training in the same human cohort to examine this further.

### One session of repeated-sprint exercise does not affect NKA α_3_, β_1_ and FXYD1 mRNA content in human muscle

We found no effect of repeated-sprint exercise on muscle NKA α_3_ mRNA content. In contrast, high-intensity exercise of substantially longer duration, relative to our protocol (40-55 min vs. 2 min), promoted α_3_ mRNA transcription (2.2-5.0 fold) in the muscle of untrained (2, 23) and trained men (30). This suggests a large exercise volume may be required to accumulate α_3_ mRNA transcripts in human muscle. In these latter studies, the α_3_ mRNA content was induced immediately post, but not 3 h following, exercise (2, 23, 30), whereas muscle biopsies were obtained after the post-exercise recovery treatment (~22 min post exercise) in the present study. This raises the possibility that the mRNA content of this gene may be induced transiently and soon after the onset of muscle activity. Another potential explanation for the present result might be associated with the large inter-individual variability (approximately 8-fold, Fig. 2C) in resting muscle α_3_ mRNA content between our participants, which may have predisposed them to different mRNA responses to exercise.

Our observation of unaltered muscle β_1_ mRNA content after one session of sprint-interval exercise is consistent with previous measurements with high temporal resolution (biopsies at 0, 1, 3, 5 and 24 h post exercise) in fibre-type heterogeneous human skeletal muscle samples after various modes of exercise (9, 11, 30, 34). Conversely, other studies of healthy men reported elevated (2.4-2.8 fold) β_1_ mRNA content after continuous, moderate-intensity cycling (23) or isolated knee-extensor exercise (10). These inconsistent human results underline that other factors than increases in the mRNA content may be of greater importance for upregulation of β_1_ protein abundance after weeks of training (41). Alternatively, our measurement time point may have reflected a transient state of mRNA with little or no effect on long-term mRNA transcript accumulation, translation and/or protein synthesis. This is underlined by the complexity of mRNA content regulation, which entails splicing, polyadenylation, mRNA export, translation and decay (40). As such, our single time-point measurement of mRNA content is a limitation of the current study, and future work should be conducted to monitor fluctuations in mRNA after repeated exercise sessions performed over several weeks with those of the corresponding protein to assess the link between mRNA and protein changes in response to exercise training.

Another novel finding was that a single session of repeated-sprint exercise did not affect FXYD1 mRNA content in human muscle. This observation extends a previous observation in rat skeletal muscle, in which FXYD1 mRNA expression remained unchanged by intense exercise (8). In the present study, FXYD1 mRNA was upregulated in some individuals by CWI. This increase coincided with substantial elevations in Sp1 mRNA content in the same individuals, indicating that changes in redox homeostasis could be important for increases in FXYD1 mRNA content in response to post-exercise CWI in human skeletal muscle. More research is required to further evaluate how exercise may affect FXYD1 mRNA in human muscle and to identify the cellular stressors involved.

### Conclusions and perspectives

This study revealed that a single session of repeated-sprint exercise increased the mRNA content of NKA α_1_ and β_3_, and decreased that of NKA β_2_. These responses were temporally associated with an increase in HIF-1α mRNA, suggesting exercise-induced muscle hypoxia may play a role in these changes. Post-exercise cold-water immersion (CWI) upregulated the mRNA content of NKA α_2_ independently of exercise alone. This effect of CWI was likely a result of modulation of muscle redox homeostasis and/or Sp1 activation. Muscle FXYD1 mRNA content was not altered by a session of sprint-interval exercise in humans. Together, these results provide novel insights into how the NKA isoforms are transcriptionally regulated by exercise and CWI in humans. Given the transcriptional information provided by the present study, future work should examine the effect of repeated bouts of post-exercise CWI performed over several weeks on NKA function and –isoform protein content.

## Additional information

## Competing interests

The authors have no conflict of interest that relates to the content of this article.

## Author contributions

Exercise testing, training and mRNA analyses were performed at Institute of Sport, Exercise and Active Living (ISEAL), Victoria University, Melbourne, VIC 3011. All authors contributed to drafting and critically revising of this manuscript. DC, JRB and DJB contributed to the conception and design of experiments. DC, JRB, DJB and JK contributed to the collection and analysis of data. DC RMM, JRB, JB, MJM and DJB contributed to data interpretation. All authors approved the final version of this manuscript.

## Funding

This study was partly funded by Exercise and Sports Science Australia (ESSA, no. ASSRG2011). DC was supported by an International Postgraduate Research Scholarship from Victoria University, Melbourne, VIC 3011, Australia.

## Acknowledgements

We thank the participants for their contribution to the study.

## References

1. Clausen, T. (2013) Quantification of Na+,K+ pumps and their transport rate in skeletal muscle: functional significance. The Journal of general physiology 142, 327–345

2. Murphy, K. T., Snow, R. J., Petersen, A. C., Murphy, R. M., Mollica, J., Lee, J. S., Garnham, A. P., Aughey, R. J., Leppik, J. A., Medved, I., Cameron-Smith, D., and McKenna, M. J. (2004) Intense exercise up-regulates Na+,K+-ATPase isoform mRNA, but not protein expression in human skeletal muscle. The Journal of physiology 556, 507–519

3. Thomassen, M., Murphy, R. M., and Bangsbo, J. (2013) Fibre type-specific change in FXYD1 phosphorylation during acute intense exercise in humans. The Journal of physiology 591, 1523–1533

4. Kristensen, M., and Juel, C. (2010) Na+,K+-ATPase Na+ affinity in rat skeletal muscle fiber types. J Membr Biol 234, 35–45

5. Harmer, A. R., Ruell, P. A., McKenna, M. J., Chisholm, D. J., Hunter, S. K., Thom, J. M., Morris, N. R., and Flack, J. R. (2006) Effects of sprint training on extrarenal potassium regulation with intense exercise in Type 1 diabetes. Journal of applied physiology 100, 26–34

6. Thomassen, M., Christensen, P. M., Gunnarsson, T. P., Nybo, L., and Bangsbo, J. (2010) Effect of 2-wk intensified training and inactivity on muscle Na+-K+ pump expression, phospholemman (FXYD1) phosphorylation, and performance in soccer players. Journal of applied physiology 108, 898–905

7. Perry, C. G., Lally, J., Holloway, G. P., Heigenhauser, G. J., Bonen, A., and Spriet, L. L. (2010) Repeated transient mRNA bursts precede increases in transcriptional and mitochondrial proteins during training in human skeletal muscle. The Journal of physiology 588, 4795–4810

8. Rasmussen, M. K., Juel, C., and Nordsborg, N. B. (2011) Exercise-induced regulation of muscular Na+-K+ pump, FXYD1, and NHE1 mRNA and protein expression: importance of training status, intensity, and muscle type. American journal of physiology. Regulatory, integrative and comparative physiology 300, R1209–1220

9. Nordsborg, N., Bangsbo, J., and Pilegaard, H. (2003) Effect of high-intensity training on exercise-induced gene expression specific to ion homeostasis and metabolism. Journal of applied physiology 95, 1201–1206

10. Nordsborg, N., Thomassen, M., Lundby, C., Pilegaard, H., and Bangsbo, J. (2005) Contraction-induced increases in Na+-K+-ATPase mRNA levels in human skeletal muscle are not amplified by activation of additional muscle mass. American journal of physiology. Regulatory, integrative and comparative physiology 289, R84–91

11. Nordsborg, N. B., Kusuhara, K., Hellsten, Y., Lyngby, S., Lundby, C., Madsen, K., and Pilegaard, H. (2010) Contraction-induced changes in skeletal muscle Na(+), K(+) pump mRNA expression - importance of exercise intensity and Ca(2+)-mediated signalling. Acta physiologica 198, 487–498

12. Petersen, A. C., Murphy, K. T., Snow, R. J., Leppik, J. A., Aughey, R. J., Garnham, A. P., Cameron-Smith, D., and McKenna, M. J. (2005) Depressed Na+-K+-ATPase activity in skeletal muscle at fatigue is correlated with increased Na+-K+-ATPase mRNA expression following intense exercise. American journal of physiology. Regulatory, integrative and comparative physiology 289, R266–274

13. Wendt, C. H., Gick, G., Sharma, R., Zhuang, Y., Deng, W., and Ingbar, D. H. (2000) Up-regulation of Na,K-ATPase beta 1 transcription by hyperoxia is mediated by SP1/SP3 binding. The Journal of biological chemistry 275, 41396–41404

14. Wang, G., Kawakami, K., and Gick, G. (2007) Regulation of Na,K-ATPase alpha1 subunit gene transcription in response to low K(+): role of CRE/ATF- and GC box-binding proteins. Journal of cellular physiology 213, 167–176

15. Ryu, H., Lee, J., Zaman, K., Kubilis, J., Ferrante, R. J., Ross, B. D., Neve, R., and Ratan, R. R. (2003) Sp1 and Sp3 are oxidative stress-inducible, antideath transcription factors in cortical neurons. The Journal of neuroscience: the official journal of the Society for Neuroscience 23, 3597–3606

16. Lafleur, V. N., Richard, S., and Richard, D. E. (2014) Transcriptional repression of hypoxia-inducible factor-1 (HIF-1) by the protein arginine methyltransferase PRMT1. Molecular biology of the cell 25, 925–935

17. Versey, N. G., Halson, S. L., and Dawson, B. T. (2013) Water immersion recovery for athletes: effect on exercise performance and practical recommendations. Sports medicine 43, 1101–1130

18. Roberts, L. A., Raastad, T., Markworth, J. F., Figueiredo, V. C., Egner, I. M., Shield, A., Cameron-Smith, D., Coombes, J. S., and Peake, J. M. (2015) Post-exercise cold water immersion attenuates acute anabolic signalling and long-term adaptations in muscle to strength training. The Journal of physiology

19. Broatch, J. R., Petersen, A., and Bishop, D. J. (2014) Postexercise cold water immersion benefits are not greater than the placebo effect. Medicine and science in sports and exercise 46, 2139–2147

20. Gregson, W., Allan, R., Holden, S., Phibbs, P., Doran, D., Campbell, I., Waldron, S., Joo, C. H., and Morton, J. P. (2013) Postexercise cold-water immersion does not attenuate muscle glycogen resynthesis. Medicine and science in sports and exercise 45, 1174–1181

21. Juel, C., Hostrup, M., and Bangsbo, J. (2015) The effect of exercise and beta2-adrenergic stimulation on glutathionylation and function of the Na,K-ATPase in human skeletal muscle. Physiological reports 3

22. Selman, C., Grune, T., Stolzing, A., Jakstadt, M., McLaren, J. S., and Speakman, J. R. (2002) The consequences of acute cold exposure on protein oxidation and proteasome activity in short-tailed field voles, microtus agrestis. Free radical biology & medicine 33, 259–265

23. Murphy, K. T., Medved, I., Brown, M. J., Cameron-Smith, D., and McKenna, M. J. (2008) Antioxidant treatment with N-acetylcysteine regulates mammalian skeletal muscle Na+-K+-ATPase alpha gene expression during repeated contractions. Experimental physiology 93, 1239–1248

24. Friedmann-Bette, B., Schwartz, F. R., Eckhardt, H., Billeter, R., Bonaterra, G., and Kinscherf, R. (2012) Similar changes of gene expression in human skeletal muscle after resistance exercise and multiple fine needle biopsies. Journal of applied physiology 112, 289–295

25. Vandesompele, J., De Preter, K., Pattyn, F., Poppe, B., Van Roy, N., De Paepe, A., and Speleman, F. (2002) Accurate normalization of real-time quantitative RT-PCR data by geometric averaging of multiple internal control genes. Genome biology 3, Research0034

26. Livak, K. J., and Schmittgen, T. D. (2001) Analysis of relative gene expression data using real-time quantitative PCR and the 2(-Delta Delta C(T)) Method. Methods (San Diego, Calif.) 25, 402–408

27. Cohen, J. (1988) Statistical power analysis for the behavioral sciences. 2nd edition

28. Galuska, D., Kotova, O., Barres, R., Chibalina, D., Benziane, B., and Chibalin, A. V. (2009) Altered expression and insulin-induced trafficking of Na+-K+-ATPase in rat skeletal muscle: effects of high-fat diet and exercise. American Journal of Physiology-Endocrinology and Metabolism 297, E38–E49

29. Ikeda, K., Nagano, K., and Kawakami, K. (1993) Anomalous interaction of Sp1 and specific binding of an E-box-binding protein with the regulatory elements of the Na,K-ATPase alpha 2 subunit gene promoter. Eur J Biochem 218, 195–204

30. Aughey, R. J., Murphy, K. T., Clark, S. A., Garnham, A. P., Snow, R. J., Cameron-Smith, D., Hawley, J. A., and McKenna, M. J. (2007) Muscle Na+-K+-ATPase activity and isoform adaptations to intense interval exercise and training in well-trained athletes. Journal of applied physiology 103, 39–47

31. Nordsborg, N. B., Kusuhara, K., Hellsten, Y., Lyngby, S., Lundby, C., Madsen, K., and Pilegaard, H. (2010) Contraction-induced changes in skeletal muscle Na plus, K plus pump mRNA expression - importance of exercise intensity and Ca2+-mediated signalling. Acta physiologica 198, 487–498

32. Mahoney, D. J., Parise, G., Melov, S., Safdar, A., and Tarnopolsky, M. A. (2005) Analysis of global mRNA expression in human skeletal muscle during recovery from endurance exercise. FASEB journal: official publication of the Federation of American Societies for Experimental Biology 19, 1498–1500

33. Kumar, A. P., and Butler, A. P. (1999) Enhanced Sp1 DNA-binding activity in murine keratinocyte cell lines and epidermal tumors. Cancer letters 137, 159–165

34. Murphy, K. T., Petersen, A. C., Goodman, C., Gong, X., Leppik, J. A., Garnham, A. P., Cameron-Smith, D., Snow, R. J., and McKenna, M. J. (2006) Prolonged submaximal exercise induces isoform-specific Na+-K+-ATPase mRNA and protein responses in human skeletal muscle. American journal of physiology. Regulatory, integrative and comparative physiology 290, R414–424

35. Murphy, K. T., Aughey, R. J., Petersen, A. C., Clark, S. A., Goodman, C., Hawley, J. A., Cameron-Smith, D., Snow, R. J., and McKenna, M. J. (2007) Effects of endurance training status and sex differences on Na+,K+-pump mRNA expression, content and maximal activity in human skeletal muscle. Acta physiologica 189, 259–269

36. Roberts, L. A., Muthalib, M., Stanley, J., Lichtwark, G., Nosaka, K., Coombes, J. S., and Peake, J. M. (2015) Effects of cold water immersion and active recovery on hemodynamics and recovery of muscle strength following resistance exercise. American journal of physiology. Regulatory, integrative and comparative physiology 309, R389–398

37. Clanton, T. L. (2007) Hypoxia-induced reactive oxygen species formation in skeletal muscle. Journal of applied physiology 102, 2379–2388

38. Yin, W., Yin, F. Z., Shen, W. X., Cai, B. C., and Hua, Z. C. (2008) Requirement of hydrogen peroxide and Sp1 in the stimulation of Na,K-ATPase by low potassium in MDCK epithelial cells. The international journal of biochemistry & cell biology 40, 942–953

39. Wenger, R. H. (2000) Mammalian oxygen sensing, signalling and gene regulation. Journal of Experimental Biology 203, 1253–1263

40. Garneau, N. L., Wilusz, J., and Wilusz, C. J. (2007) The highways and byways of mRNA decay. Nature reviews. Molecular cell biology 8, 113–126

41. Mohr, M., Krustrup, P., Nielsen, J. J., Nybo, L., Rasmussen, M. K., Juel, C., and Bangsbo, J. (2007) Effect of two different intense training regimens on skeletal muscle ion transport proteins and fatigue development. American journal of physiology. Regulatory, integrative and comparative physiology 292, R1594–1602

